# Investigation of 3D Printed Bioresorbable Vascular Scaffold Crimping Behavior

**DOI:** 10.1101/2023.10.26.564253

**Authors:** Caralyn P. Collins, Junqing Leng, Rao Fu, Yonghui Ding, Guillermo Ameer, Cheng Sun

## Abstract

The rise in additive manufacturing (AM) offers myriad opportunities for 3D-printed polymeric vascular scaffolds, such as customization and on-the-spot manufacturing, *in vivo* biodegradation, incorporation of drugs to prevent restenosis, and visibility under X-ray. To maximize these benefits, informed scaffold design is critical. Polymeric bioresorbable vascular scaffolds (BVS) must undergo significant deformation prior to implantation in a diameter-reduction process known as crimping which enables minimally invasive surgery. Understanding the behavior of vascular scaffolds in this step provides twofold benefits: first, it ensures the BVS is able to accommodate stresses occurring during this process to prevent failure, and further, it provides information on the radial strength of the BVS, a key metric to understanding its post-implant performance in the artery. To capitalize on the fast manufacturing speed AM provides, a low time cost solution for understanding scaffold performance during this step is necessary. Through simulation of the BVS crimping process in ABAQUS using experimentally obtained bulk material properties, we have developed a qualitative analysis tool which is capable of accurately comparing relative performance trends of varying BVS designs during crimping in a fraction of the time of experimental testing, thereby assisting in the integration of informed design into the additive manufacturing process.

## 1. Introduction

Both peripheral and coronary artery disease remain prevalent health issues in modern society, affecting an estimated 8-12 million and 18.2 million Americans respectively.^[1–3]^ Vascular stenting or scaffolding has become a commonplace solution for alleviating the blockage of blood vessels which oftentimes leads to these diseases. Even with this technique, restenosis, or the re-closing of the blood vessel post-implantation, has occurred at rates as high of 36.3 percent in bare metal stents and 23.1 percent in early drug eluting stents.^[4]^ While the movement in recent years from bare metal to drug-eluting stents signals improvements in widely available stenting solutions and suggests the potential for reduction in restenosis rates, commercially available stents are still traditionally relegated to metallic materials both due to required mechanical properties to withstand the deployment process and provide proper arterial support.

While metallic materials remain popular for the aforementioned reasons, they are unable to address several key concerns with the stenting process. The use of metallic materials without drug elution capabilities can cause oxidative imbalance and additional stress on the body post-vascularization.^[5]^ Additionally, once implanted, metallic stents remain in the body permanently, posing issues of stent migration in some cases up to a year post-implantation.^[6]^ In response to the limitations of metal-based stents, polymer-based bioresorbable vascular scaffolds (BVS) have become widely researched. These polymeric stents can be made of materials which are antioxidant and naturally degrade over time in the body.^[7]^ However, to date, commercial offerings of BVS remain limited, with one of the most popular options, Abbott Science’s Absorb™, being recalled and withdrawn off the market in 2017, and an improved bioresorbable vascular scaffold model from Abbott recently beginning its clinical testing.^[3, 8]^ While still in the early stages of commercial relevance, interest in bioresorbable vascular scaffolds in research spheres remains high. To improve upon existing BVS designs and improve feasibility in commercial spaces, focus has been placed in recent years on improvements in and characterization of mechanical properties of BVSs,^[9, 10]^ fabrication techniques,^[7, 11–13]^ and both biocompatibility and bioresorbability of devices.^[14–16]^

In recent years, molding, braiding, laser cutting, and additive manufacturing have all proven to be popular stent and BVS manufacturing techniques.^[17]^ Between manufacturing methods, approaches to stent or BVS design can differ vastly, given widely varying constraints, materials, and capabilities between methods. This is also true within additive manufacturing itself, wherein drastically different materials are utilized in different 3D printing techniques. Though there are multiple types of additive manufacturing which are popular in BVS production, it has become evident that 3D printing via stereolithography is an especially viable technique for producing BVS with fine features in a time-efficient manner. Its high resolution, compatibility with biocompatible materials, short fabrication times, and potential for customization and on-the-spot manufacturing offer significant benefit over numerous other techniques. Micro-continuous liquid interface production (µCLIP), the most recent evolution of stereolithography technologies, utilizes an oxygen-permeable membrane. Use of this membrane creates an oxygen-enriched deadzone at the bottom of the resin bath in which the liquid resin is prevented from curing via oxygen inhibition of free radicals, allowing for continuous replenishment of the liquid resin at the focal plane where the image is projected. Cross-sectional images can then be projected onto the focal plane of the resin bath, and the resin is cured in a layer-by-layer fashion. The use of µCLIP methods has previously enabled the 3D printing of high-resolution BVSs with 7.1 x 7.1 micron lateral resolution and 5 micron layer resolution in under thirteen minutes.^[18]^ These BVSs have high homogeneity and surface quality, enabled by the continuous nature of the printing.

Both previously and in the present study, a highly biocompatible class of citrate-based biomaterials with FDA-approved clinical applications was utilized in BVS fabrication.^[13]^ Through this method, the authors have in the past been able to manufacture mechanically reinforced composite and X-ray visible radiopaque versions of these BVS.^[18, 19]^

Given the prevalence of coronary and peripheral artery disease, it follows that much literature exists to characterize the manufacturing and *in vivo* performance of stents. This literature is spread across the wide variety of manufacturing techniques which lend themselves to stent and BVS design. However, with the increasing relevance of polymeric materials like those used in stereolithography as opposed to more traditional metallic materials, re-evaluation of performance of the BVS throughout its entire lifespan between manufacturing and deployment becomes critical. Between its manufacturing and deployment, the BVS undergoes significant amounts of deformation. First, the BVS must be compressed onto a balloon catheter via a diameter-reduction process known as crimping. This is crucial to the assembly of the BVS system for deployment, via a minimally invasive surgery. During deployment, the BVS must then be expanded back to the diameter of the vessel via inflation of the balloon catheter. In metallic stent lifespans, the performance of the stent during crimping and deployment was viewed more as a given, garnering study more as an investigation into material modeling and nuances of stent design. However, BVSs made of polymeric materials behave completely differently when undergoing large deformation associated with the crimping and deployment steps. This can be particularly challenging for the fine struts of BVS often less than 100 μm in thickness, to maintain structural integrity though the large deformation.

When accounting for polymeric vascular scaffold structural integrity during the crimping and development steps, evaluating performance by rapid design and testing iteration is by far the most common methodology. As additive manufacturing boasts high manufacturing speeds as one of its main advantages, it would be intuitive that harnessing this to quickly produce and test BVSs would be highly effective. However, this methodology relies on blind designing without ever concretely considering the material-property-process relationships. Accordingly, it often results in large amounts of wasted time and resources. In order to more decisively investigate the material-property-process relationships, a different method is necessary. In utilizing simulation to quantitatively demonstrate design performance during the crimping step, the blind nature of this design is eliminated. Hence, the use of a computer model not only offers additional information, but also potentially allows for the development of a more optimal design in a fraction of the time.

Considering their benefit to the overall design process for stents, several simulations have already been developed to investigate stent performance during crimping. McGrath et. al in 2014 developed a crimping simulation for a Nitinol stent in ABAQUS/Explicit which included a dodecagonal crimping assembly with a stent placed centrally within the crimping plates. This system was capable of analyzing radial forces the stent encountered during its deformation.^[20]^ Following this work, Shanahan, Tofail, & Tiernan developed a model in ABAQUS/Standard with both an eight-plate an octagonal crimping assembly to simulate crimping of a viscoelastic stent with each type of crimping assembly and both one and two-step analyses. This work demonstrated the viability of ABAQUS/Standard in yielding accurate results for radial forces comparatively between simulation and experiment.^[21]^ Similarly, Schiavone, Qiu, & Zhao in 2017 analyzed the crimping and expansion responses of metallic and polymeric commercial stents via ABAQUS/Explicit. This simulation involved the use of a crimping assembly, stent, and a balloon onto which the stent was crimped.^[9, 22]^ Qiu, Song, and Zhao in 2018 followed the 2017 study by analyzing simulated crimping and expansion responses of popular polymer BVS, again in ABAQUS/Explicit.^[23]^ However, not much work has yet been done to analyze the effects of crimping on 3D printed stents, and research in this area is relatively limited. Cabrera et. al in 2017 performed parallel plate crushing and crimping analysis on a 3D-printed self-expandable stent for potential heart valve replacement using fusion deposition modeling (FDM) method. Although the surface quality of the 3D printed stents is yet to be further improved, this proof-of-concept study nevertheless represents the first crimping simulation on 3D printed stents with biodegradation capacity.^[24]^

In this study, we developed a simulation model to provide accurate and more time-effective analysis of 3D printed BVS performance, using realistic BVS geometry and materials properties utilized for animal study. Similar to the published literature, we focused on the crimping process as it is well researched, easily validated, and the first time in its lifespan during which the BVS undergoes significant deformation. These BVS were composed of an in-house bioresorbable “B-ink,” and tensile testing was done to obtain mechanical properties for simulation. Using the simulation model, the performance of BVSs of varying designs and critical feature dimension were quantitatively evaluated and compared. The subject BVS designs were further 3D printed and their radial force profiles as a function of BVS diameter are measured experimentally, which provide the means to validate the simulation model. While both simulation and experimental measurements were able to discern qualitative differences in radial force between BVS designs, the simulation revealed localized stress concentrations that may lead to structural failure during crimping, which can be beneficial for rapid design iteration and optimization.

## 2. Setup & Results

### 2.1 Setup

#### 2.1.1 Experimental Setup

In order to obtain polymer bulk mechanical properties, dog bone samples for tensile testing were printed using an home-made µCLIP printer. BVSs were also printed for radial force profile validation. In this process, computer-aided design (CAD) files for the designs were initially created in SolidWorks. Each design was then converted into an STL file and sliced into 5-micron layers to be fed into the digital light projector during the printing process. These “layer” images were loaded into the printer as PNG files and projected onto the photopolymerizable ink via use of the µCLIP printer (**Figure 1a**). The samples were printed via the coordinated Z-axis motion of the building platform and the sequential projections of each of the layer images. The 3D printed samples were postprocessed in order to ensure full crosslinking and maximize mechanical strength. This printing and postprocessing procedure is further discussed in Section 4.2.

**Figure 1.**
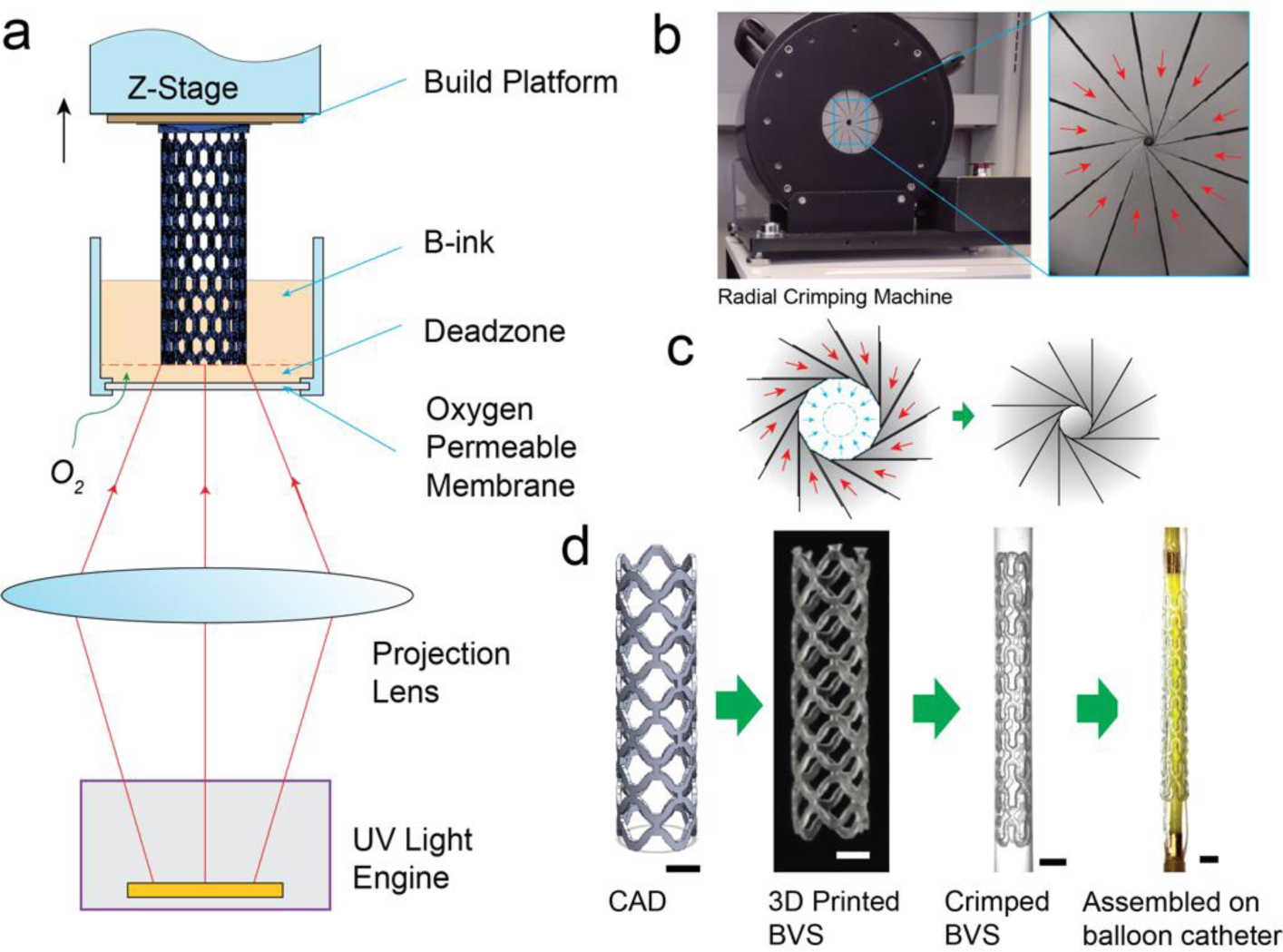
Overview of 3D printed BVS fabrication and crimping processes. a) Schematic illustration of home-made µCLIP system for 3D print BVSs and tensile testing samples. b) Radial force testing mechanism image with indication of plate motion during crimping. c) Diagram of crimping mechanism operation. d) Diagram of steps leading to BVS assembly for deployment, from CAD model to 3D printing, diameter reduction via crimping, and finally, assembly on a balloon catheter for deployment. Crimped BVS shown is held within a Teflon tube to preserve its shape. All scale bars are 1 mm.

For BVS samples, radial force analysis was completed via the use of a radial force testing mechanism (RX650, Machine Solutions, Inc.). This mechanism is shown in **Figure 1b**. Samples were placed in the center of the twelve-plate assembly, and their diameters were reduced via the translation of plates along each other to produce an inward compression on the BVS (**Figure 1c**). For in vivo testing, this process can be utilized to prepare the BVS on a catheter assembly for deployment (**Figure 1d**). Further detail on this process is given in Section 4.3.

#### 2.1.2 Simulation Setup

For efficient simulation of radial forces, a model was set up in ABAQUS/Standard. The origin of a cylindrical coordinate system was defined at the center of the BVS, with the *z*-axis oriented along the center axis of the BVS and the *r-θ* plane parallel to its end facets (**Figure 2a**). The node located at the geometric center was fixed in the theta and z-directions in order to prevent rigid body motion and ensure the stent deformed during simulation. Based upon existing stent crimping literature, the BVS was placed in the center of a twelve-plate rigid surface matching the experimental crimping system (**Figure 2b**). The rigid surface was fixed in all non-radial directions (**Figure 2c**), and in order to crimp the BVS, a displacement in the radial direction was applied to a reference point constrained to the corner nodes of each plate.^[20, 21]^ The outer diameters of the 3D printed BVSs were measured experimentally using the radial crimping machine, deeming 0.3 N radial force to be the point which corresponded to BVS diameter to ensure a reliable plate-BVS contact. BVS CAD models were then modified to reflect respective printed BVS dimensions and yield results which more precisely matched experimental testing. After crimping, experimental BVS thicknesses were also measured using ImageJ analysis of images taken via optical microscope, and the thicknesses of the simulated BVSs were adjusted to match those of experimental BVSs where necessary. The BVSs were meshed with C3D8I elements to avoid element hourglassing, and the plates were meshed with one SFM3D4R element per plate, similarly to existing literature.^[22]^ Global element sizing was determined based on a convergence test of radial force and maximum Von Mises stress values for one representative BVS design. Based on this test, four elements through the thickness resulted in a converged solution. Hence, each BVS was meshed with four elements through the thickness in the same or smaller global sizing. A coefficient of friction of 0.2 was used for all simulations.

**Figure 2.**
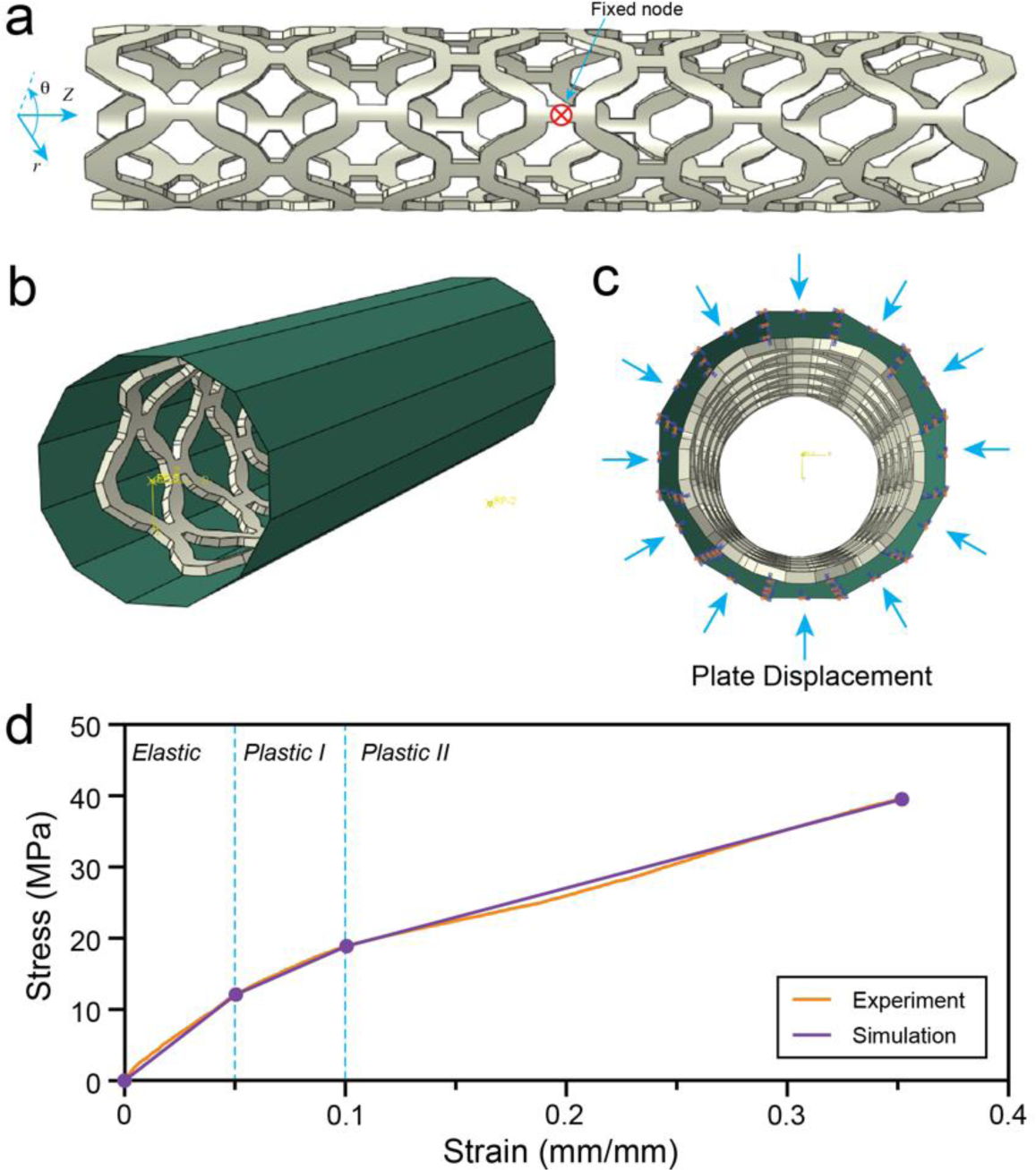
Simulation model setup in ABAQUS/Standard. a) 3D model of a representative BVS design. Fixed node boundary condition for BVS. b) Image of simulation assembly. c) Plate boundary conditions, with direction of plate displacement indirectly applied via reference point notated via blue arrows. d) Experimentally measured material properties and material property approximation used in simulation.

Each wave BVS simulation had a plate assembly diameter of 2.70 mm, and each plate was set to displace by up to 0.45 mm, resulting in a total plate assembly displacement of up to 0.9 mm. Meanwhile, the arrowhead BVS simulation had a plate assembly diameter of 3.516 mm, and each plate was set to displace by up to 0.375 mm, resulting in a total plate displacement of up to 0.75 mm. Analysis for each simulation was only utilized for radial displacements for which the BVS designs had not yet reached the ultimate tensile strength (UTS) defined by the material properties. When stresses reached the material’s UTS, failure of the scaffold was presumed, and simulation results were ignored beyond this point.

#### 2.1.3 ABAQUS material profile development

As stated in Section 2.1.1, small specimens fabricated via the same µCLIP printer are used to capture the characteristic performance of both material and processing. However, due to the small specimen size, the displacement of gage region could no longer be accurately measured with general methods; rather, only the total displacement of the whole specimen could be accurately measured. In order to avoid overestimation of sample strain due to the contribution of the two shoulder regions of the tensile specimen, prior work by Chi-tang Li and Neal Langley was followed to establish an analytical model that could extract the gage region displacement from the direct measurement of total displacement.^[25]^ For this testing, the dog bones were printed in three different gage region lengths so that the true bulk Young’s modulus of the material could be accessed (**Figure S1**). The strain of the samples of varying lengths was then calibrated to obtain an accurate averaged strain-stress curve from the specimen from which mechanical properties could be extracted.

Because this simulation was intended to provide quick analysis of BVS performance, calibrated tensile results were converted into an elastic-plastic model for use in ABAQUS (**Figure 2d**). While cyclic loading tests suggest that the material may in reality have a more complicated profile, since the simulation at hand was focused on qualitative design analysis, this profile was chosen for its low computational cost and compatibility with readily available equipment for determining mechanical properties. Yield strength was estimated via visual analysis of the calibrated curve; it was defined as the point at which the curve appeared to no longer be linear. Slope of the region between the initialization of the test and the yield point was utilized as the Young’s modulus. From this point, plastic strain was calculated based upon the chosen yield strength and modulus. One point was selected as an additional plastic strain point to bisect the plastic region based upon the apparent change in slope in the plastic region of the curve, and the end point for the curve was selected as the final stress-strain point inputted into ABAQUS, giving the profile shown in Figure 2d. The exact mechanical properties imported into ABAQUS for the material are presented in **Table 1**.

**Table 1.** Mechanical properties used in ABAQUS simulation for mPDC material.

| Young's Modulus<br>(MPa) | Poisson Ratio | Plastic Strain Points<br>(mm mm<sup>-1</sup>) | Yield Stress Points<br>(MPa) |
| --- | --- | --- | --- |
| 239.84 | 0.45 | 0 | 12.09 |
|  |  | 0.02 | 18.90 |
|  |  | 0.19 | 39.52 |

#### 2.1.4 BVS Designs

In both experiment and simulation, four BVS designs were tested. These designs can be categorized into two groups: an “arrowhead” design with 125.8 µm nominal thickness which was previously developed to maximize stent rigidity in providing arterial support,^[7]^ and a “wave BVS” design in which there are two opposing “waves” of struts, mimicking traditional metal stent geometries, which was previously developed in part due to its ability to distribute stresses over a large surface area to better accommodate the crimping process.^[19]^ The wave BVS was specifically designed to be crimped down to an outer diameter of 1.1 mm for assembly onto a catheter system and deployment *in vivo*. Within the wave BVS design group, three radial thicknesses of stent were tested: 80 µm, 110 µm, and 155 µm. The solid models of the aforementioned designs are shown in **Figure 3a**, and the corresponding printed BVSs are shown in **Figure 3b**. All designs were created using commercial Computer Aided Design (CAD) software (SolidWorks) and exported in STL format for 3D printing and STEP format for simulation.

**Figure 3.**
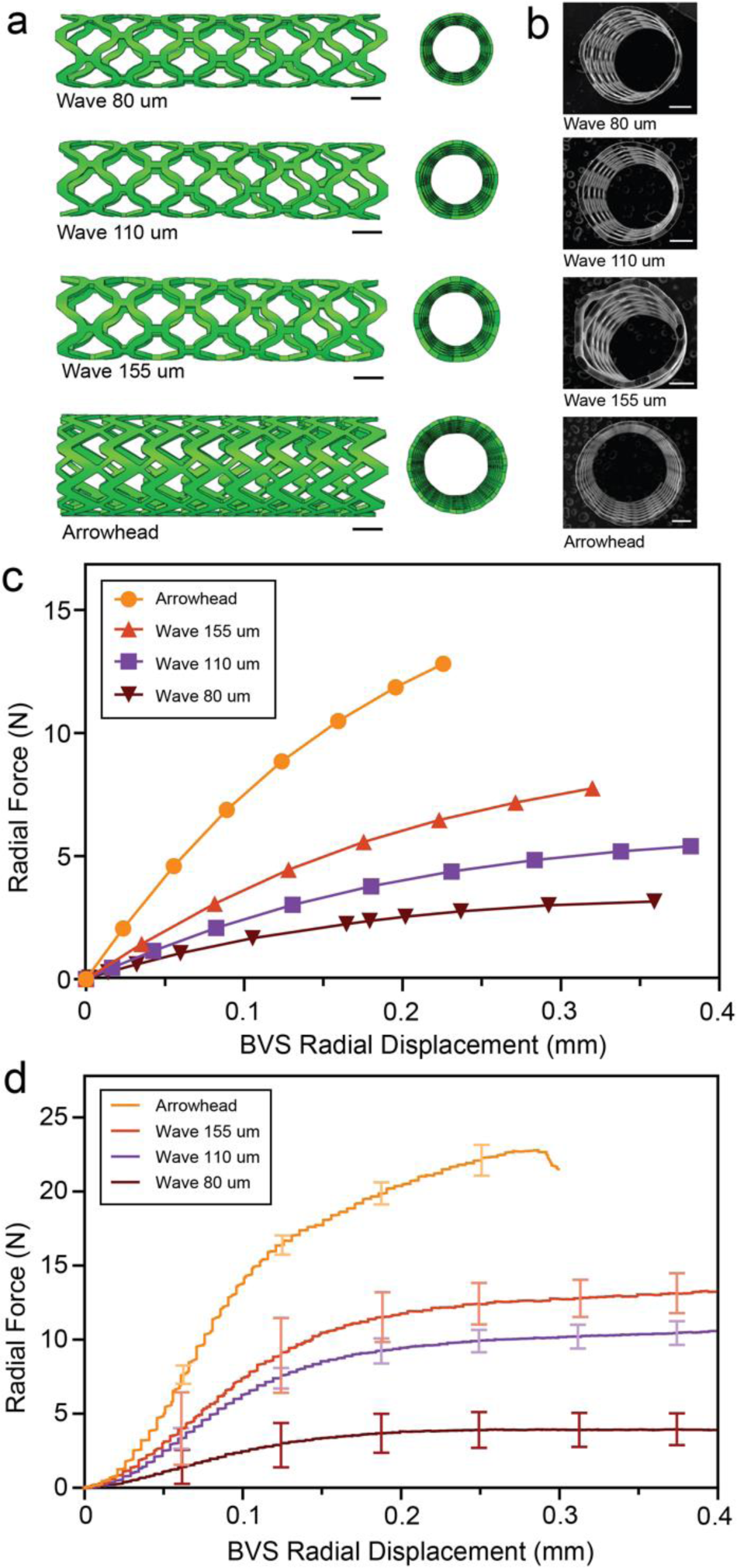
Simulated and experimentally measured crimping behavior of 4 candidate BVS designs. a) Uncrimped BVS designs in ABAQUS. b) Scanning electron microscopy (SEM) images of printed stents corresponding to the designs shown in a). c) Numerical simulation of the radial force-displacement curve during the crimping process. d) Experimentally measured radial force-displacement curve during the crimping process. All scale bars in figure are 1mm.

### 2.2 BVS Simulation Validation via Radial Force Analysis

Upon the completion of model setup with CAD models and experimentally measured materials properties, simulation was conducted to yield the force-displacement curves for the three wave BVS designs and the arrowhead design shown in Figure 3a. Amounts of crimping in this simulation (and hence final simulated BVS diameter as shown in **Figure 3c**) were determined by simulation setup as described in Section 2.1.2. Plate start position and displacement was held constant across wave BVS designs and was adjusted for the arrowhead BVS design. Final BVS diameter varied according to the initial outer diameter of each BVS design and the amount of crimping each BVS could withstand without exceeding its UTS.

The simulation results shown in Figure 3c suggest that the arrowhead BVS with a nominal thickness of 125.8 µm had the greatest radial strength, reaching a maximum radial force of 12.81 N. In contrast, the wave BVS design with a larger radial thickness of 155 µm exhibited reduced radial strength, reaching a maximum radial force of 7.76 N. The wave BVS designs with radial thickness of 110 µm and 80 µm show further reduced maximum radial force of 5.19 N and 3.16 N, respectively. Overall, the wave BVS designs exhibit rigidity positively correlated with their radial thickness while being outperformed in rigidity by the arrowhead BVS. Purely based on radial strength compared to BVS thickness, the arrowhead BVS seems to be a good candidate for *in vivo* applications. However, the stresses it encounters during crimping, which will be discussed in detail in Section 2.3, significantly limit its performance.

We further conducted experimental tests to qualitatively validate the simulation model (Section 2.1.1 & Section 4.2-4.3). This experimental BVS radial force testing yielded force-displacement curves that are directly comparable to simulation results (**Figure 3d**). In analyzing this data, the arrowhead BVS once more demonstrated high rigidity, with its maximum radial force reaching 23.13 N. At the same time, this design encountered a significant drop in radial force after around 0.6 mm of total crimping (corresponding to 0.3 mm of radial BVS displacement), indicating the occurrence of structural failure during the crimping process. Meanwhile, the 155 µm thick wave BVS was the next most rigid, with a maximum radial force of 13.61 N. The 155 µm thick wave design did not experience any significant drops in radial force for the analyzed diameters, indicating its ability to withstand crimping. However, based upon its comparable strut thickness to the recalled Absorb™ BVS, this design is not necessarily feasible clinically, as negative impact of BVSs with this thickness on vessel health has been demonstrated.^[3, 26]^ The 110 µm thick wave BVS was also able to withstand crimping well, seeing no significant drops in radial force for the analyzed diameters. Its radial strength was still relatively close to the force range of the 155 µm thick wave BVS, reaching a maximum of 10.95 N. The 80 µm thick wave BVS, while in large part able to withstand crimping (only one sample in this group showed failure during experimental testing), showed a large reduction in radial strength from the 110 µm thick wave BVS. Its maximum radial force was 4.29 N, just under half that of the 110 µm thick wave BVS design.

The wave BVS designs were specifically created to accommodate the BVS crimping process, allowing ample room for struts to fold during crimping without causing failure (as discussed in Section 2.1.4). This was consistent with the ability of the BVS to be crimped 0.6 by mm or more overall, or 0.3 mm radially, without failure both in simulation and experiment. Different BVS thicknesses for this design were utilized to investigate the role radial thickness plays regarding BVS strength and ability to distribute stresses. Increasing BVS thickness in the radial dimension increased radial strength both in simulation and experiment. Meanwhile, the arrowhead BVS design was created much earlier for sole purpose of maximizing its rigidity without considering the crimping process. This is true both in simulation and experiment, as the arrowhead design had much higher radial strength than any of the wave BVS designs but was unable to handle as much crimping as the wave designs without typically encountering some sort of failure (further discussed in Section 2.3).

The qualitative agreements between the experiments and simulation thus validate the accuracy of the simulation model. Notably, simulation model is capable of differentiating BVS responses of the arrowhead BVS and wave BVS designs, as well as the wave BVS designs of varying thickness. Notably, this simulation was sensitive enough to predict differences in radial force profiles of BVSs which varied in radial thickness by as little as 30 µm. The remaining quantitative differences between simulation and experiment are likely attributed to two prevailing factors. First, the printed BVS geometries are slightly different than those utilized in simulation. When utilizing μCLIP, corners tend to be smoother than they are in design, resulting in a different distribution of stresses than is seen in theoretical design. Additionally, mechanical properties in simulation do not align perfectly with those experienced in simulation. While the two profiles match well, cyclic testing of the material in experiment revealed a viscoplastic mechanical profile, in contrast to the elastic-plastic one that was utilized for this simulation. The elastic-plastic profile was kept to allow use in ABAQUS/Standard and provide a quick framework for material importation into ABAQUS from experimental testing. Further, the properties utilized are tensile, whereas the BVS here is undergoing compression. Both of these factors are discussed further in Section 2.5.

### 2.3 BVS Von Mises Stress Analysis

While radial force analysis validates the simulation, further investigation of mechanical BVS deformation is necessary in order to truly gain an understanding of how design details such as surface area in high-stress regions, shape of strut bending points, and radial thickness impact BVS crimping performance both on a local and global scale. In order to better understand these relationships, the spatial distribution of Von Mises stresses was further simulated (**Figure 4**).

**Figure 4.**
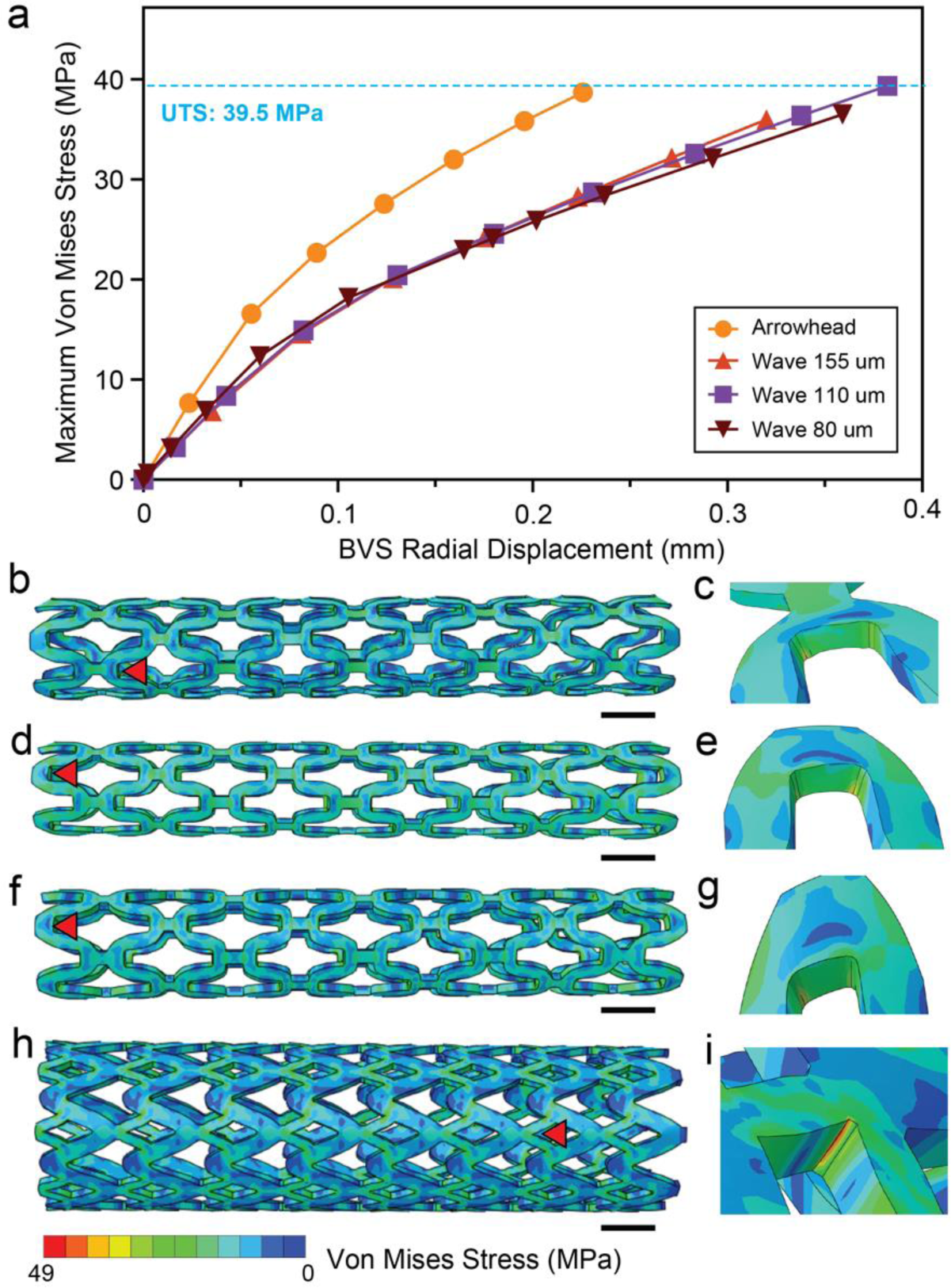
Simulation of Von Mises stresses varying in gradation and magnitude across different designs and strut thicknesses. a) Von Mises stresses over course of simulation for point which has the maximum stress integration point at the end of crimping. b) BVS stress profile at most crimped condition for 80 µm thick wave BVS design. c) Local stress profile for 80 µm thick wave BVS surrounding maximum stress integration point at the end of crimping. d) BVS stress profile at most crimped condition for 110 µm thick wave BVS design. e) Local stress profile for 110 µm thick wave BVS surrounding maximum stress integration point at the end of crimping. f) BVS stress profile at most crimped condition for 155 µm thick wave BVS design. g) Local stress profile for 155 µm thick wave BVS surrounding maximum stress integration point at the end of crimping. h) BVS stress profile at most crimped condition for arrowhead BVS design. i) Local stress profile for arrowhead BVS surrounding maximum stress integration point at the end of crimping. Red triangles in b)-h) indicate the unit ring for which the corresponding local stress profiles are shown. All scale bars in figure are 1 mm.

Regions which have the highest stress concentrations indicate areas in which the BVS may fail during crimping. This information was first captured via analysis of the maximum Von Mises stress at integration points correlating to the maximum visualization nodal Von Mises stresses (**Figure 4a**). Because of the choice in element used for the simulated BVS mesh, the nodal Von Mises stress is different from that at the integration points.

As can be seen in Figure 4a, the arrowhead BVS design approached the material’s ultimate tensile strength at a much smaller amount of crimping than any of the wave BVS designs. Here, the arrowhead BVS design approaches the material UTS at a radial displacement value of less than 0.25 mm, corresponding to a total BVS diameter reduction of 0.5 mm, whereas the wave BVS designs do not begin to reach the material UTS until a radial displacement of more than 0.3 mm, corresponding to an overall BVS diameter reduction of 0.6 mm. The Von Mises stresses of the wave BVS designs, however, were much less differentiated than their radial strengths. At the end of crimping, the 155 μm thick wave BVS had the highest stress of the three wave BVS thicknesses relative to the amount of radial displacement of the BVS, followed by the 110 μm thick wave BVS and the 80 μm thick wave BVS, respectively. Hence, at the end of crimping, thickness did positively correlate with a higher Mises stress for the same amount of crimping as other BVS designs. However, throughout, all three of the wave BVS designs exhibit similar maximum stress curves throughout the simulation, indicating that thickness had very limited impact on the maximum Mises stresses of the wave BVSs throughout the process as a whole.

Following this analysis, gradations of stresses throughout and across BVS designs were further investigated. Global stress profiles were obtained for the 80 µm thick wave BVS (**Figure 4b**), the 110 µm thick wave BVS (**Figure 4d**), the 155 µm thick wave BVS (**Figure 4f**), and the arrowhead BVS (**Figure 4h**) respectively to better show stress variation and distribution between designs. The wave BVS designs exhibited similar stress distributions to each other globally.

Although the magnitudes of stresses may vary locally between the designs, the wave BVSs distribute stresses similarly regardless of their thicknesses. This relatively constant distribution throughout the scaffold was expected, since these BVS were originally designed to be able to be crimped 0.4-0.5 mm total (0.2-0.25 mm radially) beyond the amount they were crimped in this study. Meanwhile, the arrowhead design showed steeper gradations in stress near its small connecting struts. Overarchingly, the arrowhead design had larger stresses at these points than was the case for the wave BVS designs. This is consistent with the expected behavior for this BVS due to its design for rigidity.

Local stress profiles around the points of maximum stress were also obtained for the 80 µm thick wave BVS (**Figure 4c**), the 110 µm thick wave BVS (**Figure 4e**), the 155 µm thick wave BVS (**Figure 4g**), and the arrowhead BVS (**Figure 4i**) to elucidate the roles of surface area and shape of the region about which struts were bending during crimping. Here, the gradations in stress around the locations of maximum stress were once again similar between the three wave BVS designs; there were two stress concentrations in the interior of the region about which the BVS struts bent. This design was created to have two points about which BVS struts bent in order to ensure the BVS could be crimped to a diameter of 1.1 mm, so this follows the expected trend.

However, the location of maximum stress did change between designs, with the location for the 80 µm thick BVS being an interior unit ring, while the location for the 110 µm thick and 155 µm thick wave BVS designs was at an end unit ring. Meanwhile, the arrowhead BVS showed a singular stress concentration near to its maximum stress location. This is expected given that it was only designed to have one region about which the struts bent during crimping.

Since the arrowhead BVS utilized in this study was one which was initially created without consideration of the BVS crimping process, this design was used as a baseline for analysis. Notably, stresses in the arrowhead designs were much higher at small crimping amounts than those in the wave BVS designs. In simulation, it was seen that the BVS reached its Von Mises stress close to the ultimate tensile stress after approximately a half of a millimeter of overall crimping or 0.25 mm of radial displacement (Figure 4a). For the arrowhead BVS design, this was a reliable predictor of failure during experimental BVS crimping; in experiment, when the arrowhead BVS was crimped, buckling was observed in each trial (n=5), with a drop in radial force on the BVS occurring at an average diameter of 2.52 mm, after around 0.6 mm of total crimping, corresponding to a radial dispacement of 0.3 mm (Figure 3c). This suggests that this design is not sufficient for allowing the BVS to be crimped to a required diameter for in situ use of around 1 mm.

Buckling and BVS failure were not widely observed in this experiment for the wave BVS designs (observed only in one 80 µm thick wave BVS design). Across the BVSs, varying gradation of stresses was observed between arrowhead and wave BVS designs. In the case of the arrowhead BVS designs, gradations in stress (Figure 4h-i) were much steeper than those of the wave BVS designs. Further, high stress regions were much more concentrated than in the wave BVS designs (Figure 4b-g). Correspondingly, the wave BVS designs seemed to distribute stress concentrations across larger regions than the arrowhead BVS designs. This suggests that the wave designs are indeed better at withstanding the crimping process without approaching any sort of failure limit, and further suggests that they would much better serve *in vivo* applications where significant amounts of crimping prior to deployment are necessary.

### 2.4 BVS Plastic Strain Analysis

Beyond analysis of Von Mises stresses, equivalent plastic strains (PEEQ) provide valuable information about which regions of the BVS deform plastically during crimping. This provides additional information on the regions of the BVS which potentially require strengthening for clinical application as well as once again providing additional design information on the relationship between the design on a global and local scale and intricacies of BVS deformation during the crimping process beyond what is obtainable solely through analysis of radial strength. In order to further elucidate this relationship, qualitative trends in plastic strain were analyzed via comparison of global and local equivalent plastic strain distributions between BVSs. The equivalent plastic strain was globally visualized for each of the 80 µm thick wave (**Figure 5a**), 110 µm thick wave (**Figure 5c**), 155 µm thick wave (**Figure 5e**), and arrowhead BVS (**Figure 5g**) designs to allow for qualitative comparison of plastic strains in various regions of the BVS between each design. The 80 µm thick wave BVS exhibited significant plastic strain on the “connecting” struts (struts in line with the length of the BVS) between the opposing wave-like struts. This seems to be correlated to the thickness of the stent, as the 110 µm thick wave and 155 µm thick wave BVS designs did not show as much PEEQ in these regions. Considering both this plastic strain and the bending of the ends of the BVS for the 80 µm thick wave BVS, it appears that this BVS may not have sufficient radial strength to both withstand the crimping process and expand back to its normal shape. This mirrors experimental testing, as during crimping, buckling was observed in one of the six 80 µm thick BVS crimped. Other than in the connecting struts, the wave BVS designs overall exhibited similar profiles of plastic strain. The thicker 155 µm thick wave BVS exhibited smaller magnitudes of plastic strain than the 110 µm and 80 µm thick wave designs. This suggests that plastic strain was negatively correlated with BVS radial thickness.

**Figure 5.**
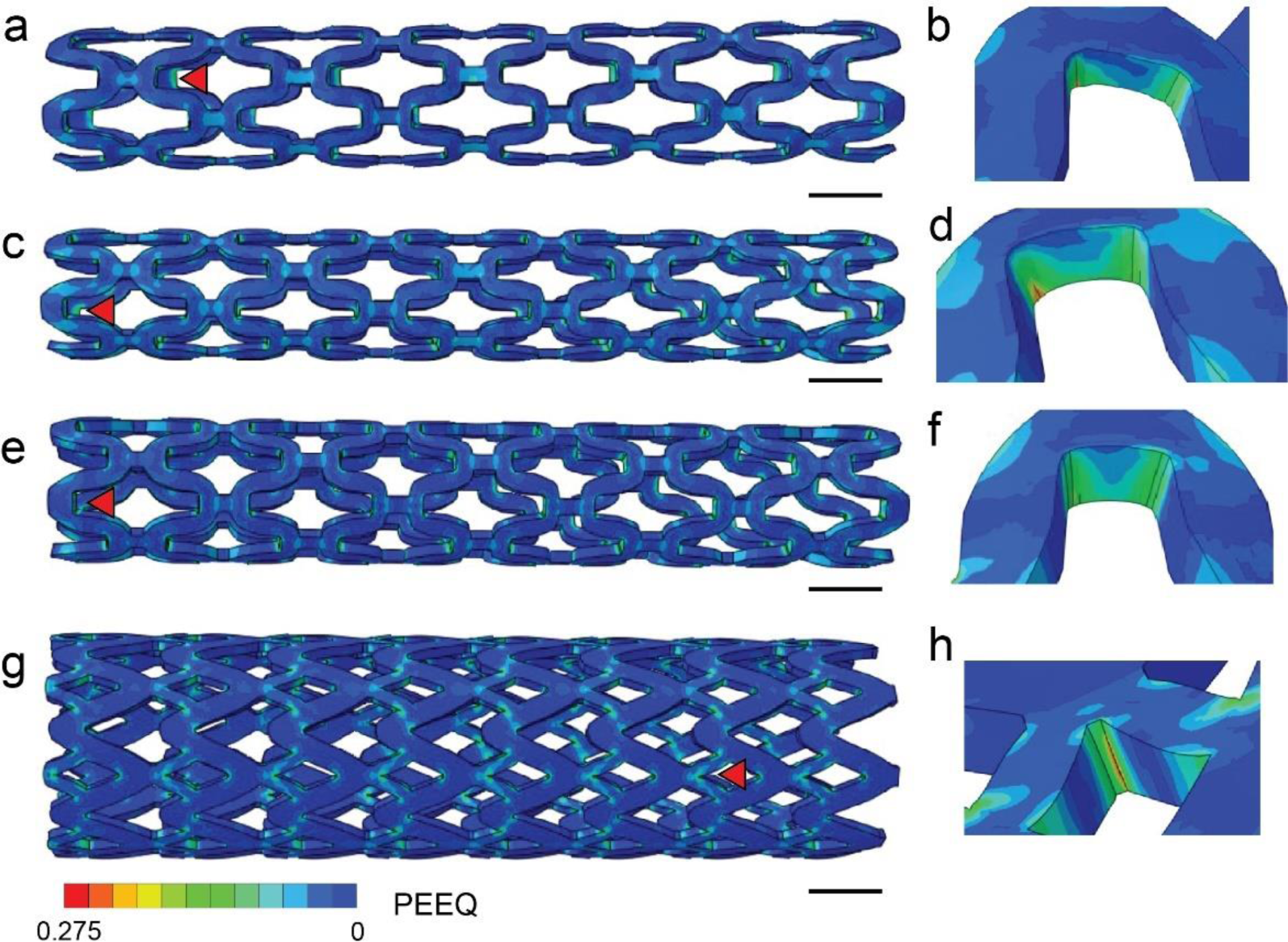
Simulation of equivalent plastic strains (PEEQ) varying according to both BVS thickness and design type. a) Global equivalent plastic strain visualization for 80 µm thick wave BVS. b) Local equivalent plastic strain visualization for 80 µm thick wave BVS surrounding point of maximum PEEQ at end of crimping. c) Global equivalent plastic strain visualization for 110 µm thick wave BVS. d) Local equivalent plastic strain visualization for 110 µm thick wave BVS surrounding point of maximum PEEQ at end of crimping. e) Global equivalent plastic strain visualization for 155 µm thick wave BVS. f) Local equivalent plastic strain visualization for 155 µm thick wave BVS surrounding point of maximum PEEQ at end of crimping. g) Global equivalent plastic strain visualization for arrowhead BVS. h) Local equivalent plastic strain visualization for arrowhead BVS surrounding point of maximum PEEQ at end of crimping. All strains are given in mm mm^-1^. Red triangles in a-g indicate unit ring for which the corresponding local strain profiles are shown. All scale bars in figure are 1 mm.

The equivalent plastic strain was also analyzed locally in the region around the point of maximum equivalent plastic strain for each of the 80 µm thick wave (**Figure 5b**), 110 µm thick wave (**Figure 5d**), 155 µm thick wave (**Figure 5f**), and arrowhead BVS (**Figure 5h**) designs to better inform design of the region about which struts were bending during crimping. The shape of the of equivalent plastic strain profile appeared similar around the region of maximum strain between the 80 µm, 110 µm, and 155 µm thick wave BVS designs. However, the arrowhead BVS had a significantly different shape of strain profile, with a singular steep strain concentration. When comparing the plastic strain of the arrowhead BVS to that of the wave BVS designs, the regions of plastic strain are much more heavily concentrated in the arrowhead BVS design (Figure 5g-h). When the relative surface areas of the regions about which each BVS’s struts bend during crimping are compared, this follows expected trends—the arrowhead BVS, with smaller surface area about which each strut bends, exhibits greater and more concentrated plastic deformation. This again suggests that this design may not be sufficient for situations in which crimping is crucial to the BVS’s successful deployment.

### 2.5 Simulation Versus Experimental Time Cost

In analyzing results as they compare to rapid experimental testing, simulation of this process provides accurate qualitative analysis of varying BVS design in a fraction of the time required for rapid experimental testing. Once setup was completed, each simulation could be run in sub-3 hour time frames, with a combined process time of less than five hours (291 minutes) for the four simulations run here. Eight-processor parallelization was configured in ABAQUS to allow for more time-efficient simulation. Meanwhile, the experimental process resulted in a process time of over 36 hours. Hence, while rapid design iteration and experimental testing did provide a greater degree of quantitative information, simulation resulted in a roughly seven-times increase in speed of design analysis. Simulation was also able to provide qualitative information on specific behavior of each design which was experimentally unavailable. Furthermore, simulation remains a much more financially accessible tool than rapid experimental testing, costing significantly less to utilize than the required crimping mechanism and materials. To the best of the authors’ knowledge, we demonstrate here the first simulation which allows for accurate qualitative analysis of relative 3D printed BVS performances in a short timespan, thus eliminating the need for time and resource-expensive blind design iteration.

### 2.6 Comparison to Rapid Experimental Testing & Future Work

There were several noticeable differences in results between rapid experimental testing and simulation. The first of these was the quantitative difference between radial force values observed in experiment and simulation. The most likely contributor to this difference is the calculation of material properties for the material at hand. The polymer used for both tensile samples and BVSs in this study is one which is manufactured in-house. In order to gather mechanical properties, tensile specimens are printed at reduced sizes from ASTM standards and calibrated accordingly. The 3D printing of these specimens likely results in some defects which result in reduced material strength when tensile testing, which may compound with the smaller specimen size to potentially give different mechanical properties from the true bulk material.

Beyond this, cyclic loading of the samples suggests that the material is likely viscoplastic, which would require a more robust material testing process and a more computationally expensive simulation setup in order to be fully represented. Finally, mechanical properties used in this simulation were tensile, and compressive testing would likely give a more accurate representation of the material. Another potential reason for quantitative differences between radial force values observed between experiment and simulation is the differences between printed BVS geometry and simulated BVS geometry. When printed, BVS structures tend to encounter more rounded corners and smoother transitions between struts than are present in the CAD model. This could potentially result in varying stress distributions and radial forces encountered between printing and simulation.

One additional difference between rapid experimental testing and simulation was the shape of the beginning of the force-displacement curve. In simulation, the BVS was placed at the center of the plate system, and radial forces first began to occur when all plates contacted the BVS’s outer surface. However, in experiment, the BVS was placed at the bottom of the twelve-plate system, resulting in plate contact with part of the BVS in certain regions of the outer surface before all BVSs were contacted. This may contribute to the different profile shape at the beginning of the experimental force-displacement curve.

Analysis of simulated results opens the door for multiple avenues of work in the future. First, there are refinements that can potentially be made to improve simulation accuracy in analysis of material properties of 3D printed samples, potential inclusion of compressive testing results, and study of how to best represent the material at hand. Additional work can also be done to ensure the BVS crimping in experiment more precisely matches the simulation setup. Beyond that, this simulation was meant as a tool to help alleviate the need for rapid experimental testing in the design generation process. While on its own, this simulation helps reduce that need, a more effective solution would be to utilize this simulation as part of a larger system for predicting performance of 3D printed designs via machine learning. This would allow users to potentially not only enter designs into simulation to predict performance, but to be able to optimize their design in a fracture of the time using technology as an aid.

## 3. Conclusion

This work presents an ABAQUS/Standard simulation of the BVS crimping process for various 3D printed BVS geometries, using experimentally obtained material properties. We have shown that the simulation can capture qualitative differences in performance between 3D printed BVSs of varying designs, which are further validated via experimental measurements. Furthermore, simulation also provided important qualitative information about trends in stress distribution and plastic strain for varying BVS geometries beyond what is experimentally obtainable, allowing for better understanding of the potential failure modes or structural weak points in the BVS design. This simulation offers promising potential in prompting rapid design convergence toward optimal performance. This work can be further extended to modeling the balloon dilatation step to fully capture the full life cycle of the BVS, providing additional insight into *in vivo* deployment.

## 4. Methods

### 4.1 Ink Formulation

Samples were fabricated from a photopolymerizable “B-ink” previously developed by Ware et. al consisting of 75% methacrylated poly(1,12 dodecamethylene citrate)--henceforth referred to as mPDC--as monomer, 19.8% ethanol as solvent, 2.2% Irgacure 819 as the primary photoinitiator, and 3% Ethyl 4-(dimethylamino)benzoate (EDAB) as co-photoinitiator by weight, respectively.^[13]^ The citrate to diol molar ratio used for the mPDC in this study was 2:1, and the synthesis process generally followed was that used by Van Lith et. al and Ding et. al.^[7, 19]^ Irgacure 819 and EDAB were obtained from Sigma Aldrich (St. Louis, MO). Following the addition of each of these components into a glass bottle, the mixture was sonicated until homogeneous, for approximately one and a half hours. The bottle was then covered with black aluminum foil to prevent polymerization of the ink due to ambient light and was placed in a freezer for storage.

### 4.2 Sample Printing & Postprocessing

All samples for the experimental sections of this study were printed via the use of a custom μCLIP digital light projection-based system. CAD designs were completed in SolidWorks, saved as STL files, and sliced in five micron layers. These layers were then projected onto the focal plane of the resin bath via the use of a digital light projector, and cross-sectional projections were sequentially coordinated with the z-motion of the printer’s moving fabrication stage. Lateral pixel size for this system was approximately 7.1 μm, and z-resolution was 5 μm.

For postprocessing, samples were first agitated in an ethanol rinse for approximately a minute and were subsequently placed under UV flood for 2 minutes, rotated, and subjected to 2 more minutes of UV flood. Samples were then placed in a 120°C oven for 10 hours, similarly to previously reported procedures with a slight modification in oven cure time.^[13]^ Samples were then stored in phosphate-buffered saline (PBS) on a hot plate at 37°C for at least 24 hours prior to testing to mimic BVS *in vivo* conditions and allow for the collection of more situationally accurate bulk material properties.

### 4.3 Radial Force Testing

To test the radial strength of the BVSs and validate the simulation, a commercial radial force tester (RX650, Machine Solutions, Inc.) was used (Figure 1b). Plates in this system slide along each other to reduce the size of the center aperture and provide a diameter-reducing compressive force on the BVS (Figure 1b,c). A 10 lbf load cell was used for testing. The plate diameter was calibrated using the gauge pin with preselected diameters of 4 and 30 mm. The load cell was further calibrated using 5 and 10 lb weights prior to testing. After calibration, crimping ramp profiles were started at a plate diameter of 4 mm, slightly larger than the BVS outer diameters of up to 3.1 mm, and crimped the BVS down to an outer diameter of 1.1 mm. Data from these profiles was used to obtain an averaged experimental force-displacement curve for 0.8 mm of diameter change from the initial BVS diameter for each BVS geometry, corresponding to 0.4 mm of radial change, or until significant drops in radial force, indicative of BVS failure, were observed. For each BVS design, at least n=5 samples were tested.

### 4.4 Tensile Testing for Mechanical Properties

A single column tabletop model testing system (Instron 5940 Series) equipped with a 2 kN load cell was used for tensile testing. Dog bone sample dimensions were measured and input into the machine. The sample was subsequently loaded into the tensile testing grips and a tensile force was applied. Samples were tested at a displacement rate of one millimeter per second. Upon failure, the samples were removed and replaced. After testing, the stress-strain curve data points from the Instron were then converted into true stress-strain values. Young’s modulus was calculated manually via calibration of the engineering stress-strain profile obtained from tensile testing (Figure 2d). For each dog bone type, at least n=3 samples were tested.

## Supporting Information

Supporting Information is available from the Wiley Online Library or from the author.

## Acknowledgements

The authors gratefully acknowledge Dr. Henry Oliver Tenadooah Ware for his work on CAD for the BVSs used in this project. This work was supported by the National Institute of Health [grant number #R01HL141933 and R01DE030480]. Y. Ding was in part supported by the Center for Advanced Regenerative Engineering and the American Heart Association (AHA) Career Development Award [grant number 852772]. *This work made use of the EPIC facility of Northwestern University’s NUANCE Center, which has received support from the SHyNE Resource (NSF ECCS-2025633), the IIN, and Northwestern’s MRSEC program (NSF DMR-2308691)*.

## Conflict of Interest

The authors declare no known conflict of interest for this work.

## Notes

### Competing Interest Statement

The authors have declared no competing interest.

