## Supplementary material for "Investigation of 3D Printed Bioresorbable Vascular Scaffold Crimping Behavior": Figure S1

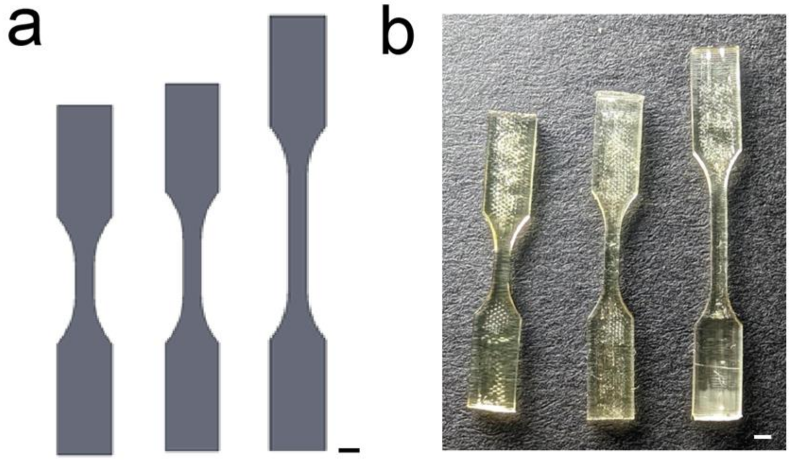


**Figure S1.** Dog bone designs for mechanical testing of mPDC. a) CAD files for dog bones with gauge regions of varying length. b) As-printed dog bones matching designs shown in a). All scale bars in figure are 1 mm.
